# Loss of E-cadherin Induces IGF1R Activation Revealing a Targetable Pathway in Invasive Lobular Breast Carcinoma

**DOI:** 10.1101/2022.02.22.480917

**Authors:** Ashuvinee Elangovan, Jagmohan Hooda, Laura Savariau, Susrutha Puthanmadhomnarayanan, Megan E. Yates, Jian Chen, Daniel D. Brown, Priscilla F. McAuliffe, Steffi Oesterreich, Jennifer M. Atkinson, Adrian V. Lee

## Abstract

No Special Type breast cancer (NST; commonly known as Invasive Ductal Carcinoma (IDC)) and Invasive Lobular Carcinoma (ILC) are the two major histological subtypes of breast cancer with significant differences in clinicopathological and molecular characteristics. The defining pathognomonic feature of ILC is loss of cellular adhesion protein, E-cadherin (*CDH1*). We have previously shown that E-cadherin functions as a negative regulator of the Insulin-like Growth Factor 1 Receptor (IGF1R) and propose that E-cadherin loss in ILC sensitizes cells to growth factor signaling which thus alters their sensitivity to growth factor signaling inhibitors and their downstream activators. To investigate this potential therapeutic vulnerability, we generated CRISPR-mediated *CDH1* knockout (*CDH1* KO) IDC cell lines (MCF7, T47D, ZR75.1) to uncover the mechanism by which loss of E-cadherin results in IGF pathway activation. *CDH1* KO cells demonstrated enhanced invasion and migration that was further elevated in response to IGF1, serum and Collagen I. *CDH1* KO cells exhibited increased sensitivity to IGF resulting in elevated downstream signaling. Despite minimal differences in membranous IGF1R levels between wildtype (WT) and *CDH1* KO cells, significantly higher ligand-receptor interaction was observed in the *CDH1* KO cells, potentially conferring enhanced downstream signaling activation. Critically, increased sensitivity to IGF1R, PI3K, Akt and MEK inhibitors was observed in *CDH1* KO cells and ILC patient-derived organoids, suggesting that these targets require further exploration in ILC treatment and that *CDH1* loss may be exploited as a biomarker of response for patient stratification.

## Introduction

Despite significant diagnostic and therapeutic advancements, breast cancer remains a leading cause of malignancy associated mortality for women worldwide. Breast cancer is a heterogenous disease that can be categorized based on its histopathology and molecular features. The major histological subtypes of breast cancer are No Special Type (NST) which is often referred to as Invasive Ductal Carcinoma (IDC: ∼80% of all cases) and Invasive Lobular Carcinoma (ILC: ∼10-15% of all cases) (1–3). Although IDC and ILC are clinically managed as the same disease, there is an increasing appreciation of their differences in terms of molecular alterations, pathology, and prognosis (4–7). The diminished prevalence of ILC consequently has resulted in this subtype being historically understudied and therefore implores the need to better characterize the disease to establish precision medicine approaches to treatment (4, 8). ILC tumors are largely characterized by their growth in a single file, discohesive manner with presence of stromal infiltration (5–7, 9). Approximately 95% of ILC cases present with loss of *CDH1*, with the majority due to genomic alterations such as loss of heterozygosity and/or truncating frameshift mutations (6, 10, 11). In addition, other molecular alterations such as *PIK3CA* activation, PTEN loss, and mutations in TBX3 and FOXA1 are detected at higher levels in ILC tumors compared to IDC tumors (4, 12). Enhanced Akt signaling has also been observed in several ILC cell line and mouse models (13).

As one of the activators of Akt, the Insulin-like Growth Factor 1 (IGF1) pathway has long been associated with breast cancer progression (14–18). IGF1 is an essential component in cellular proliferation and survival (15, 19). Binding of IGF1 to its receptor (IGF1R) or the insulin receptor (IR) stimulates autophosphorylation and activation of the PI3K and MAPK pathways among others (15, 18, 20). Due to high levels of circulating IGF1 being associated with poor prognosis in breast cancer (21), it is imperative to understand the dysregulation of the IGF1 pathway in tumors. Numerous IGF1R inhibitors have been evaluated in preclinical and clinical settings, including humanized monoclonal antibodies that downregulate the receptor and small molecule tyrosine kinase inhibitors which inhibit pathway activation (20, 22–26). Unfortunately, these trials demonstrated little to no response in the recruited broad patient populations despite promising preclinical efficacy (13, 27–31), suggesting the necessity to better stratify patients to ensure efficient validation of therapies while maximizing response metrics.

To that end, we recently reported E-cadherin as a modulator of IGF1 signaling and a potential biomarker of inhibitor response (32). Briefly, transient E-cadherin knockdown sensitized cell lines to IGF1 signaling and to IGF1R inhibition (32, 33), and we further reported that ILC tumors have higher IGF1/2 expression and pIGF1R/IR activation compared to IDC tumors (13, 33). In this present study, we have expanded upon our earlier observations by generating CRISPR-mediated *CDH1* knockout (*CDH1* KO) IDC cell lines to elucidate the role of E-cadherin individually in more detail. Here we report that loss of E-cadherin renders cells sensitive to IGF1, IGF2 and insulin signaling by increasing IGF1R receptor availability for ligand binding resulting in enhanced cell migration/invasion and contextual increases in sensitivity to IGF1R, PI3K, Akt and MEK inhibitors. Given previous evidence for E-cadherin regulation of other growth factor pathways (13, 30), we also extended our analyses to additional growth factor signaling pathways. Our findings provide insights into better understanding the mechanics of E-cadherin function as well as the potential for a targeted therapeutic intervention in ILC patient populations.

## Materials and Methods

### Cell culture

Cell lines utilized in this study were obtained from ATCC: MCF7 (RRID: CVCL_0031), T47D (RRID: CVCL_0553), ZR75.1 (RRID: CVCL_0588), MDA-MB-134-VI (RRID: CVCL_0617), MDA-MB-231 (RRID: CVCL_0062) and Asterand for SUM44PE (RRID: CVCL_3424). Cell lines were maintained in 10% fetal bovine serum (FBS; Life Technologies) supplemented media (Thermo Fisher Scientific): MDA-MB-134 in 1:1 DMEM: L-15; MCF7 and MDA-MB-231 in DMEM; and T47D and ZR75.1 in RPMI. SUM44PE was maintained in DMEM/F12 with 2% charcoal stripped serum (CSS; Life Technologies) with additional supplements as previously described (5). Cell lines were cultured for less than 6 months at a time, routinely tested to be Mycoplasma free and authenticated by the University of Arizona Genetics Core (Tucson, Arizona) by short tandem repeat DNA profiling.

### *CDH1* knockout cell line generation

CRISPR mediated knockout of *CDH1* in MCF7 and T47D cells was performed by utilizing the Gene Knockout Kit (V1) from Synthego (Redwood City, California) as previously described (31, 34). The parental cell line was used a comparator for MCF7 and T47D, referred as wildtype (WT). For ZR75.1, a doxycycline inducible Cas9 was utilized harboring either short guide RNA (sgRNA) for *CDH1* or a non-targeting control (NTC) sequence and was introduced via an adenoviral vector. For ZR75.1, a total of 3 rounds of infection and Puromycin selection was performed before single cell sorting. 8 clones from each KO and NTC cells were isolated by single cell cloning and combined to generate a pool for subsequent experiments.

### Haptotaxis, migration and invasion assays

For haptotaxis experiments, the QCM Haptotaxis Cell Migration Assay Collagen I (EMD Millipore #ECM582) kit was used as previously described (8). For Transwell migration assays, transparent 24 well PET membranes of 8µm pore size (Fisher Scientific # 08-771-21) were used. For Collagen I invasion assays, QCM Collagen Cell Invasion Assay (EMD Millipore #ECM551) was used according to manufacturer’s protocol. For the latter two assays, cells were plated at a density of 300,000 cells/well in 300µL 0.5% FBS media in the top chamber; all bottom chambers were filled with 0.5% FBS media +/- 5nM IGF1 (GroPep Bioreagents # AQU100) or full serum (10%) media. Cells were incubated at 37°C for 72 hours. Excess cells were removed from the top chambers using cotton swabs and inserts were stained with Crystal Violet (Sigma-Aldrich #C0775) before being imaged on an Olympus SZX16 dissecting microscope and quantified with ImageJ software. Quantifications were normalized to low serum WT samples and p-values calculated with one-way ANOVA.

### Receptor availability assay

Cells were seeded in 6cm plates (Fisher #08-772-E), and serum starved overnight after achieving a 70-80% confluency. Cells were then stimulated with biotinylated IGF1 (GroPep #AQU100) for 15 minutes at 4°C to reduce receptor internalization. Following PBS washes on ice, cells were treated with 2mM BS3 crosslinker (Thermo Scientific #21580) reconstituted in PBS (pH 8.0) with 6mM KCl and 10mM EGTA for 1 hour at 4°C with occasional rocking. BS3 quenching was performed with 10mM Glycine for 15 minutes and cells were harvested for subsequent immunoblotting (30).

### Dose Response assays

Cells were plated in 50µL of media at 9,000 cells/well in 2D and ULA (Corning #3474) 96-well plates. Treatments were added 24 hours post seeding in an additional 50µL of respective media. IGF1R inhibitors BMS-754807 (Selleckchem #S1124) and OSI-906 (Selleckchem #S1091), PI3K inhibitor Alpelisib (Selleckchem #S2814), Akt inhibitor MK-2206 (Selleckchem #S1078), MEK inhibitor U0126 (Selleckchem #S1102) and Fulvestrant (Selleckchem #S1191) were dissolved in DMSO with a final ≤0.5% DMSO concentration in treatments. Plates were collected at day 6 and measured by CellTiter-Glo (Promega #PR-G7573) following the manufacturer’s protocol. Cell viability values were analyzed following blank cell deductions and normalization to vehicle readings. IC50 values for viability were calculated by nonlinear regression and statistical differences evaluated using sum-of-squares Global f-test (p<0.05). Synergy was assessed using SynergyFinder (35).

### In silico analysis

Gene expression data from the Sweden Cancerome Analysis Network–Breast (SCAN-B) study was downloaded from Gene Expression Omnibus, accession GSE96058 (36). Differential gene expression between luminal A lobular (N = 265) and ductal (N = 1165) breast cancer samples was assessed using DESeq2 (37). Tumor purity for SCAN-B samples was estimated using the R package ESTIMATE (37). An FDR cut-off of 0.05 was used to identify significantly differentially expressed genes. Heatmap for genes of interest was created using the R package ComplexHeatmap. Gene set variation analysis (GSVA) was performed on log transformed gene FPKM matrices using the R package GSVA (38) with default parameter “Gaussian” for kernel selection. Gene sets of interest were obtained from MSigDB version 7.4. The GSVA enrichment scores were compared between luminal A lobular and ductal breast cancer samples using Mann-Whitney U test. Gene set enrichment analysis (GSEA version 4.1.0. Broad Institute) (39) was also conducted on normalized raw counts with default parameters and customized gene sets of interest obtained from MSigDB.

### Organoid culture

IDC organoids (IPM-BO-56 and 102) and ILC organoids (IPM-BO-30, 41, 46, 77 and 114) were established by the Institute for Precision Medicine (Pittsburgh, PA) and maintained in media as detailed in (40) with 1nM β-estradiol (Sigma-Aldrich #E8875) supplementation. For dose response assays, organoids were dissociated to single cell suspension with Trypsin and plated in 50µL of organoid media at 3,000 or 5,000 cells/well in 96-well round bottom plates (Falcon #353227) depending on the growth rate. Akt inhibitor MK2206 (Selleckchem #S1078; dissolved in DMSO) treatments were added 24 hours post seeding in an additional 50µL of organoid media. Organoids were monitored every day to ensure vehicle treated wells were growing well and remained at an appropriate density. Media was refreshed on day 6 and plates collected on day 12 with cell viability quantified by CellTiter-Glo 3D (Promega #G9681). Dose response assay analysis was performed as described above.

### Data Availability Statement

All data generated in this study are available within the article and its supplementary files. SCAN-B data analyzed in this study was obtained from Gene Expression Omnibus (GEO) at GSE96058.

## Results

### IGF pathway activity is enhanced in ILC patient samples and cell lines *in vitro*

Analysis of the publicly available Sweden Cancerome Analysis Network–Breast (SCAN-B) study allowed a direct comparison of growth factor signaling differences between luminal A IDC (n=1165) and luminal A ILC (n=265) tumors. Significantly higher IGF1 and IGF2 expression were observed in ILC tumors in addition to several other growth factor signaling related genes (**Fig 1A**). PI3K/Akt signaling was significantly higher in ILC by gene set enrichment analysis (GSEA) (**Fig 1B)** and gene set variation analysis (GSVA) (**Fig 1C**). IGF1/2 signaling activation was also higher in ILC (**Fig 1D**), strengthening our hypothesis regarding IGF pathway activation in ILC over IDC tumors.

**Figure 1:**
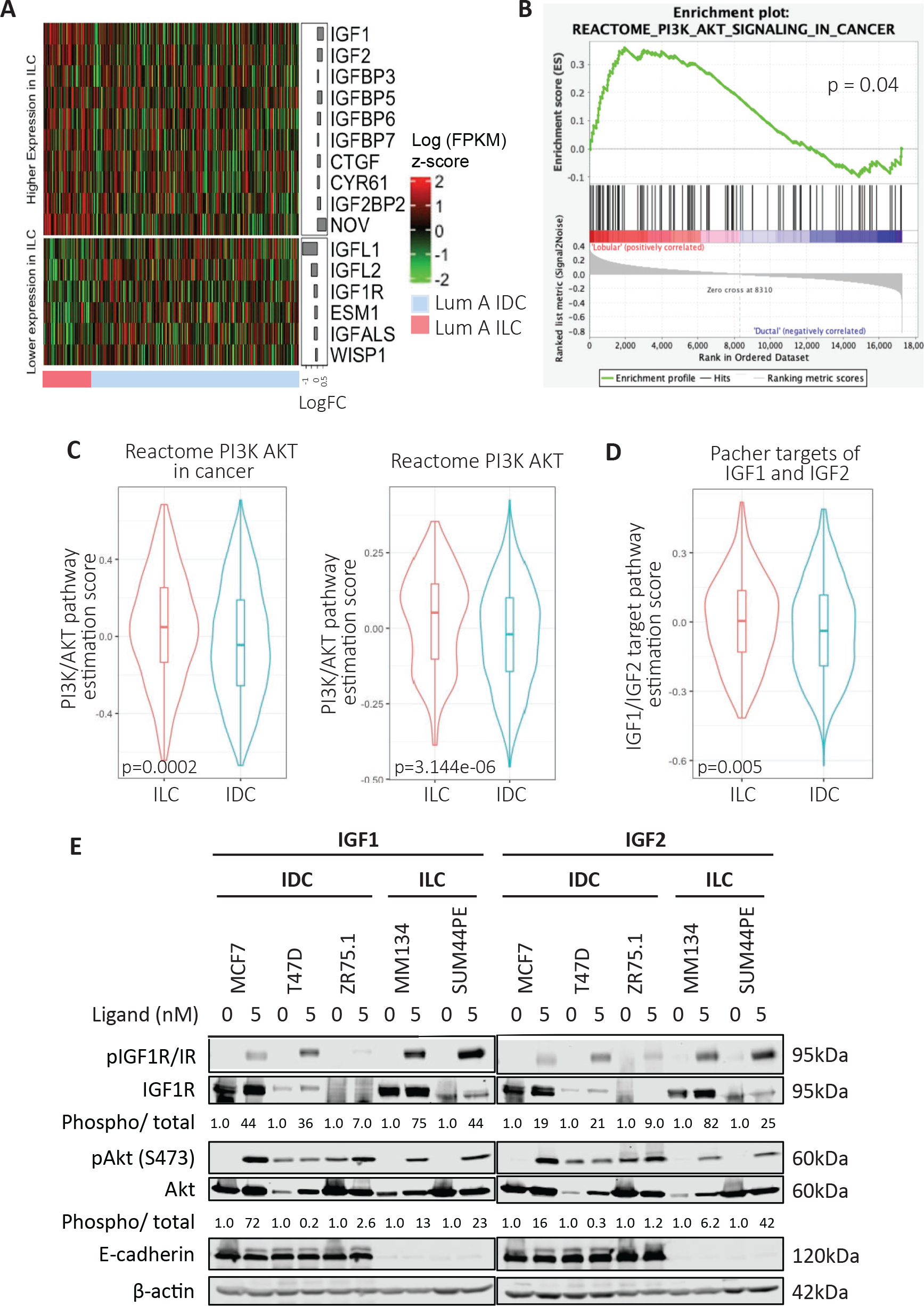
IGF pathway activity is enhanced in ILC patient samples and cell lines *in vitro*. (A) Gene expression analysis of growth factor signaling related genes comparing Luminal A IDC and ILC with significantly different expression values displayed. (B) GSEA analysis for PI3K/Akt showing significantly higher pathway activation in ILC samples obtained from the SCANB dataset. (C, D) GSVA analysis of Luminal A IDC and ILC samples using respective signature sets from MSigDB with significantly higher pathway activation in ILC. (E) IDC cells (MCF7, T47D, ZR75.1) and ILC cells (MDA-MB-134-VI, SUM44PE) were stimulated with doses of IGF1 or IGF2 (0-5nM) for 15 minutes following an overnight serum starvation. IGF1R/IR and Akt signaling was assessed by Western blotting. Phosphorylation levels of IGF1R (phospho-IGF-I Receptor β (Tyr1135/1136)/Insulin Receptor β (Tyr1150/1151)) and Akt (S473) were quantified on the LiCOr Odyssey CLx Imaging system and normalized to corresponding total protein levels and loading controls. Ligand treated sample values were further normalized to respective cell line vehicle treated samples. Representative experiment shown for all, N=2-3 for each experiment.

We assessed the differences in IGF pathway activity in IDC (MCF7, T47D, ZR75.1) and ILC (MDA-MB-134-VI, SUM44PE) cell lines. Both ILC cell lines showed higher pIGF1R/IR expression after IGF1 stimulation compared to the IDC cell lines, though the expression of total IGF1R/IR varied across cell models (**Fig 1E**). pAkt (S473) levels, as a measure of downstream pathway activation, did not always correlate with pIGF1R/IR levels. We observed Akt activation in the absence of ligands in T47D and ZR75.1 cells, while MCF7 cells demonstrated robust pAkt induction despite modest pIGF1R/IR levels. These results are not entirely surprising given the existing *PI3K* activating mutations in MCF7 and T47D (41–43), and *PTEN* loss in ZR75.1 cells (43) which may contribute to the Akt activation. We further assessed the effects of insulin and again observed that ILC cell lines demonstrated enhanced activation of pIGF1R/IR, although with this ligand ZR75.1 also showed robust activation (**Supp Fig 1A**). To test whether there was a difference between ILC and IDC cell lines in response to ligands targeting other cell surface receptors that may be regulated by E-cadherin (30, 44–46), we assessed the effects of EGF stimulation in our panel of cell lines (**Supp Fig 1B**). Little EGFR activation could be detected in ILC cell lines suggesting cellular context specific changes in growth factor signaling activation. Taken together these data support increased IGF pathway activity in ILC compared to IDC in both representative patient tumors and in a cell line models.

### *CDH1* knockout cells as a model to study the role of E-cadherin in regulating the IGF pathway

As previously reported (32, 33), and expanded herein (**Fig. 1**), E-cadherin negatively regulates IGF pathway activity. To better understand this regulation, we generated CRISPR-mediated *CDH1* knockout (*CDH1* KO) IDC cell lines of MCF7, T47D and ZR75.1. Both MCF7 and T47D demonstrate successful and complete knockout of *CDH1* while ZR75.1 *CDH1* KO cells retained some expression of E-cadherin though markedly reduced compared to WT cells (**Fig 2A, B**).

**Figure 2:**
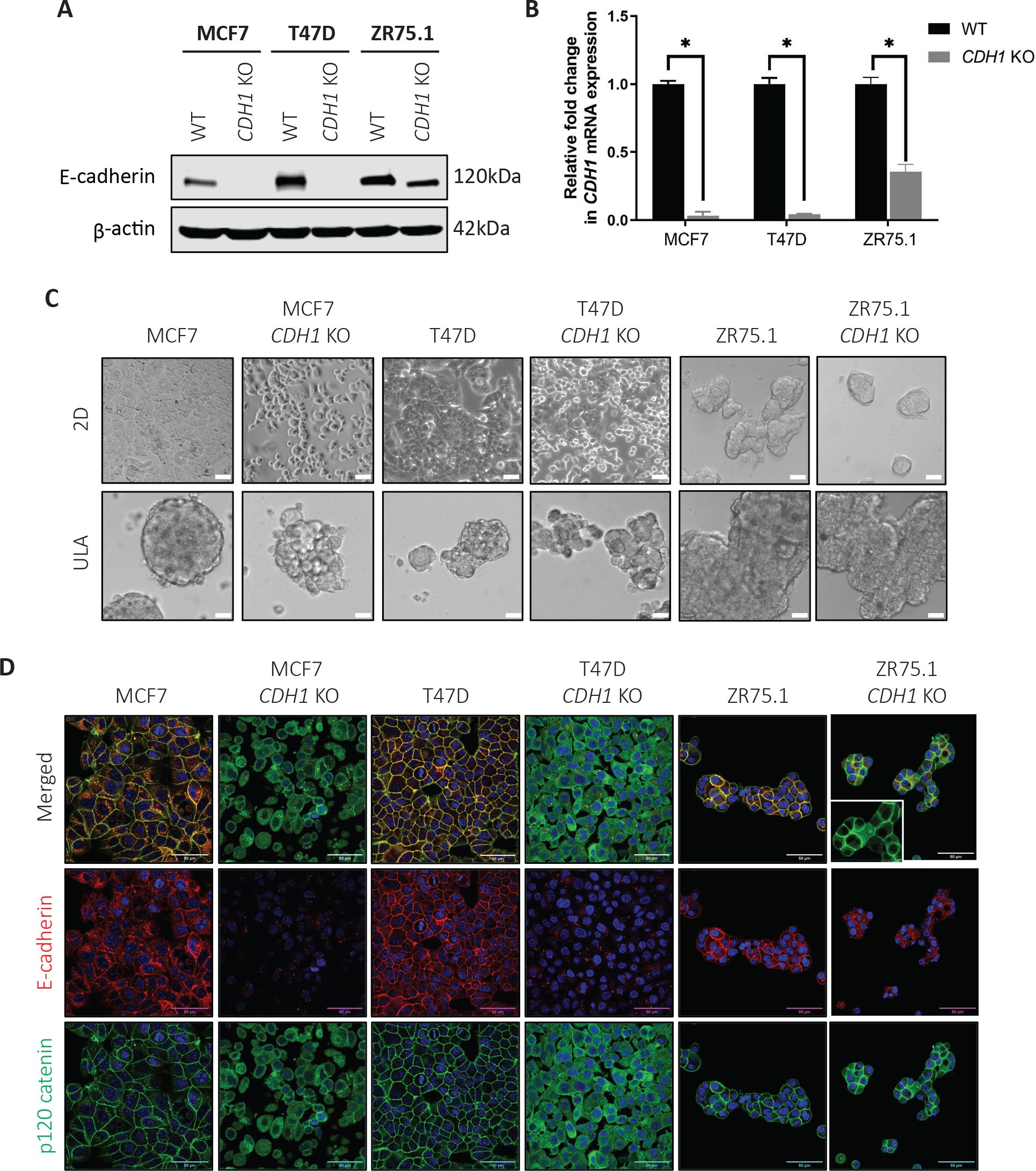
*CDH1* knockout cells as a model to study the role of E-cadherin in regulating the IGF pathway. (A) Western blotting and (B) qRT-PCR confirms reduction of E-cadherin in *CDH1* KO MCF7, T47D and ZR75.1 CRISPR cell lines compared to wildtype (WT) parental cells. Statistical differences evaluated using paired t-test (*p < 0.05). Representative experiment shown, N=2 (each with two biological and 3 technical replicates). (C) Representative brightfield images (10X magnification) of WT and *CDH1* KO cell line models plated in 2D and Ultra-low attachment (ULA) plates. (D) E-cadherin (red) and p120 catenin (green) staining of WT and *CDH1* KO cells confirms *CDH1* loss and p120 re-localization in *CDH1* KO cell models by confocal microscopy (60X objective). Inset in the ZR75.1 *CDH1* KO panel shows a zoomed in image. Scale bar: 50µm. Representative experiment shown for all, N=2-3 for each experiment.

ZR75.1 cells harbor copy number amplification of *CDH1* which we believe made complete knockout of *CDH1* in this model challenging (42, 47).

We first assessed cell morphology given the known role of E-cadherin in formation of cell-cell junctions. As shown in **Fig 2C**, MCF7 and T47D *CDH1* KO cells have a rounded morphology compared to their parental (WT) cells in 2D culture. In addition, they also demonstrated decreased cell-cell attachment as well as decreased cell-plate attachment. In ultra-low attachment (ULA) conditions, MCF7 and T47D *CDH1* KO cells appeared to form less tight, more grape-like cell clusters synonymous with ILC cell aggregates (8, 48). We did not note any obvious difference in either 2D or ULA conditions between ZR75.1 WT and *CDH1* KO cells.

Examination of E-cadherin by immunofluorescence confirmed loss in all three cell lines models (**Fig 2D**). Furthermore, similar to ILC cell lines (8, 49), p120 catenin was re-localized to the cytoplasm in the E-cadherin null *CDH1* KO cells. Consistent with our earlier observations, ZR75.1 *CDH1* KO cells showed residual E-cadherin expression, and p120 expression was visualized in both the cytoplasm and at the cell membrane.

As a function of the cadherin complex, E-cadherin stabilizes β-catenin on the cell membrane and prevents its release into the cytoplasm. The release of β-catenin results in free β-catenin being targeted for proteasomal degradation and/or its accumulation resulting in subsequent activation of Wnt signaling. We thus examined expression of β-catenin following GSK inhibitor (CHIR- 99021; Selleckchem #S2924) treatment. With vehicle treatment, both MCF7 and T47D *CDH1* KO cells exhibited little appreciable β-catenin expression, which was only elevated following 24-hour GSK inhibitor treatment (**Supp Fig 1C**), suggesting that *CDH1* KO alone does not alter β-catenin accumulation. Together, these data support the successful generation of *CDH1* KO IDC cell line models which display phenotypes consistent with loss of E-cadherin functionality including the characteristic re-localization of p120 as seen in human ILC tumors.

### Loss of E-cadherin sensitizes cells to IGF signaling pathway activation

To understand the signaling effects of E-cadherin loss, we subjected WT and *CDH1* KO cells to IGF1 stimulation. We observed enhanced sensitivity in all three *CDH1* KO cell lines by proxy of higher pIGF1R/IR and pAkt activation (**Fig 3A**). This enhanced signaling response of *CDH1* KO cells to WT cells was evident to the greatest extend upon 5nM IGF1 stimulation: 2.72-fold, 1.3- fold and 1.5-fold higher signaling in MCF7, T47D and ZR74.1 *CDH1* KO cells respectively. The pAkt elevation was not apparent in ZR75.1 cells, possibly accounted by *PTEN* loss in this model as discussed previously. We next assessed the effects of E-cadherin loss on IGF2, and insulin signaling and observed enhanced IGF1R/IR activation in MCF7 and ZR75.1 *CDH1* KO cells following ligand stimulation (**Fig 3B, C**). This enhanced signaling translated into enhanced Akt activation in the ZR75.1 *CDH1* KO cells, but not in the MCF7 *CDH1* KO cells. T47D *CDH1* KO cells showed similar sensitivity to both IGF2 and insulin when compared to WT cells, suggesting an IGF1 specific differential signaling response.

**Figure 3:**
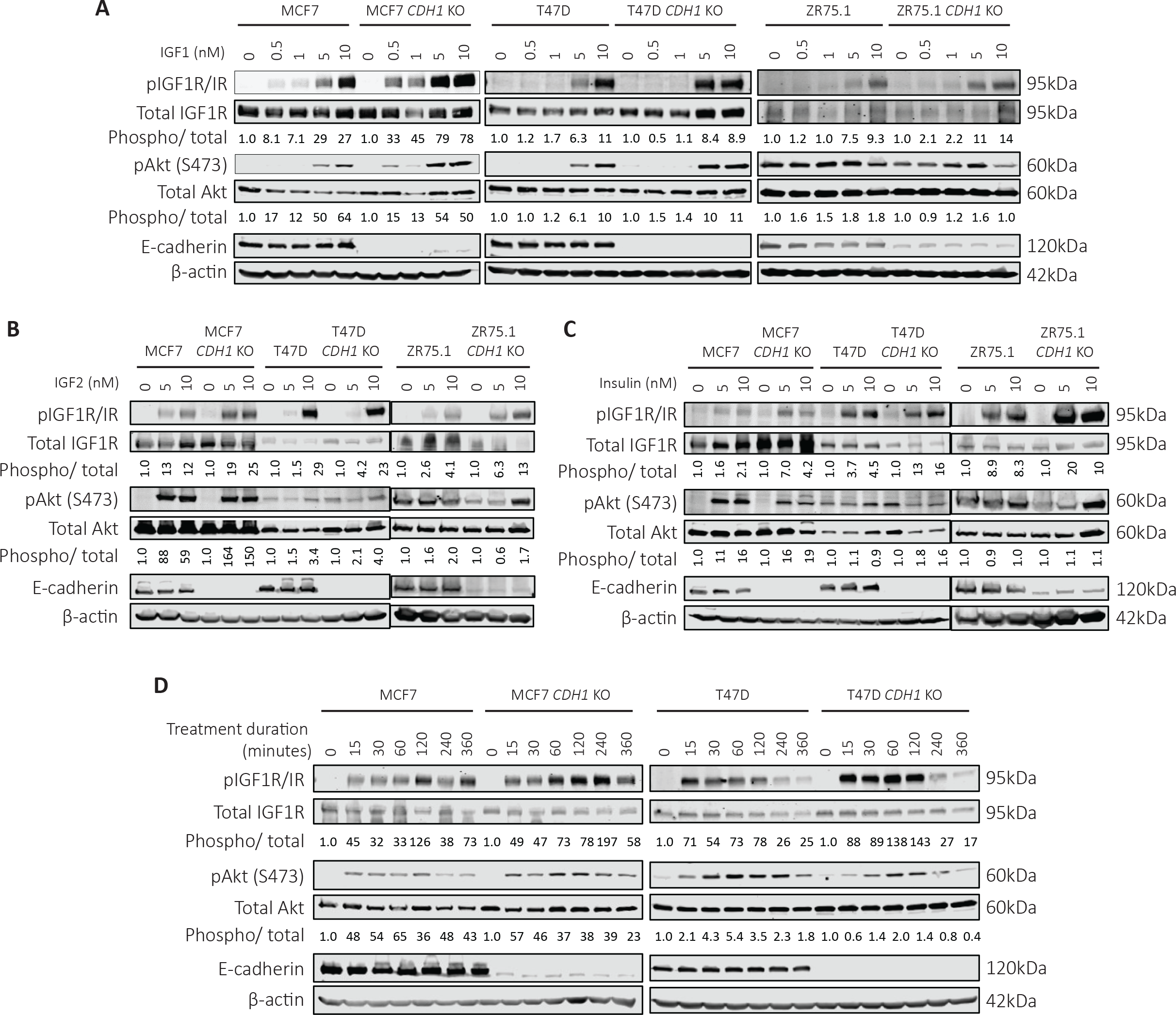
Loss of E-cadherin sensitizes cells to IGF signaling pathway activation. Cells were serum starved overnight and stimulated with (A) IGF1, (B) IGF2 or (C) Insulin (0-10nM) for 15 minutes. Cells were harvested for Western blot to assess IGF1R/IR and Akt signaling. For quantification, phosphorylated protein levels were normalized to corresponding total protein levels and loading controls. Ligand treated sample values were further normalized to respective cell line vehicle treated samples. (D) Cells were treated with 10nM IGF1 for a time course from 0-6 hours to assess the duration of signaling activity between WT and *CDH1* KO cells. Representative experiment shown for all, N=2-3 for each experiment.

To explore if our findings were related to a general growth factor sensitivity following loss of E-cadherin, we stimulated cells with EGF but did not observe any consistent differences in pEGFR Tyr1068 activation, or pAkt levels between *CDH1* KO and WT cells (**Supp Fig 2A**). We specifically examined the effects of E-cadherin deletion on FGFR activity given recent studies on the role of FGFR4 as a driver of endocrine resistance (50), particularly in ILC (51). Cells were stimulated with a cocktail of FGF ligands (FGF 1, 2, 4, 6, 8, 17, 19, 21 and 23) resulting in strong enhancement in both raw and normalized pFGFR4 levels in both MCF7 and T47D *CDH1* KO cells, despite decreased total FGFR4 levels in the *CDH1* KO cells (**Supp Fig 2B**).

Surprisingly, while pFGFR4 was elevated in *CDH1* KO cells, downstream activators did not display a difference between WT and *CDH1* KO cells. Additionally, in both MCF7 and T47D *CDH1* KO cells, we observed elevated pathway activity that was sustained for up to 4 hours in the MCF7 cells and up to 2 hours in the T47D cells (**Fig 3D**). These results were replicated for pAkt activation in the MCF7 *CDH1* KO cells but not in the T47D *CDH1* KO, warranting additional studies on the sustained receptor activity and its effects on downstream signaling response. Results obtained with ZR75.1 cells were not robust (**Supp Fig 2C**) despite multiple trials. Collectively, these results suggest that there is an enhancement of IGF1 response in IDC cell lines harboring E-cadherin loss under the conditions tested, but that this increased signaling sensitivity may not be generalizable to other cell surface receptors and is mostly applicable to the IGF1 pathway.

### Loss of E-cadherin increases IGF1R availability on the membrane to allow ligand binding

To understand why and how loss of E-cadherin leads to sensitized IGF1R signaling, we first compared the expression of IGF1R in the WT and *CDH1* KO cells. Using cell fractionation, minimal differences between the MCF7 and ZR75.1 WT and *CDH1* KO cells were noted (**Fig 4A**). Higher IGF1R expression was observed in both whole cell and membrane fractions of the T47D *CDH1* KO cells compared to WT, which was consistent with increased IGF1R mRNA (**Supp Fig 3A**). We assessed the availability of the receptor for ligand binding using a biotinylated IGF1 ligand, which was crosslinked to its receptor for subsequent complex quantification via α-biotin immunoblotting. MCF7, T47D and ZR74.1 *CDH1* KO cells showed a 2.1-fold, 4-fold and 1.7-fold higher ligand-receptor complex respectively when compared to their corresponding WT cell line controls (**Fig 4B**). This finding was stronger in the T47D *CDH1* KO cells, likely due to the higher baseline IGF1R expression in these cells. As E-cadherin has previously been reported to create a repressive EGFR complex (30), we examined for the presence of a physical interaction between IGF1R and E-cadherin on the cell membrane which potentially could physically and spatially prevent ligands from binding IGF1R. An immunoprecipitation performed for both IGF1R and E-cadherin did not reveal co-IP of either protein despite multiple attempts to optimize conditions (**Fig 4C**). Imaging of IGF1R to assess localization on the membrane showed no significant differences between the WT and *CDH1* KO cells (**Fig 4D**). Our results suggest that although no significant differences in IGF1R expression was observed between WT and *CDH1* KO cells, increased receptor availability to bind ligand likely leads to the observed elevated signaling induction upon ligand binding.

**Figure 4:**
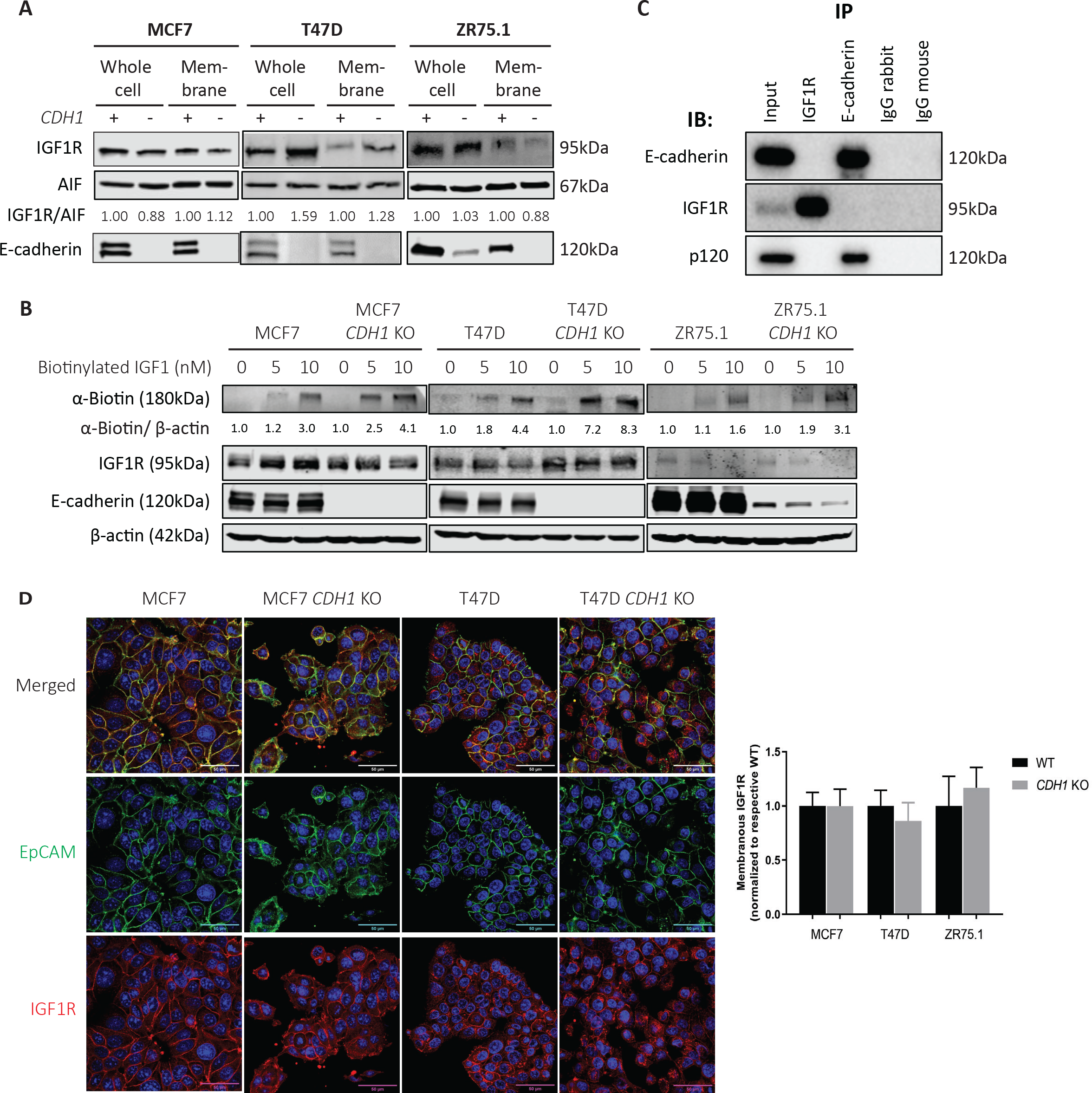
Loss of E-cadherin increases IGF1R availability on the membrane to allow ligand binding. (A) Cell fractionation assay performed on MCF7, T47D and ZR75.1 WT and *CDH1* KO cells to assess whole cell and membrane IGF1R expression levels. IGF1R bands were quantified and normalized to membrane control, AIF1. (B) Cells were stimulated with biotinylated IGF1 (0-10nM) for 10 minutes and crosslinked to assess ligand-receptor complex levels between WT and *CDH1* KO cells. (C) Immunoprecipitation of IGF1R and E-cadherin in T47D cells was assessed for a co-IP of other proteins, with p-120 catenin assessed as a known interactor of E-cadherin. (D) Cell lines were dual stained for EpCAM (green) and IGF1R (red) and imaged by confocal microscopy at a 60X objective. Scale bar: 50µm. 12 images for each cell line were quantified and graphed. Representative experiment shown for all, N=2-3 for each experiment.

### Loss of E-cadherin enhances anoikis resistance and increases cell survival

We previously showed increased anchorage independent growth in ILC cells compared to IDC (8). In IDC cells with *CDH1* KO, we similarly found an increase in ULA growth compared to WT cells in the MCF7 and T47D models (31) as has also been seen by others (52). ZR75.1 WT and *CDH1* KO cells did not show any significant differences in either growth condition (**Supp Fig 4A**). To explore the previously observed ability for MCF7 and T47D *CDH1* KO cells to thrive in ULA conditions, we examined anoikis resistance using an Annexin V and Propidium Iodide staining technique followed by flow cytometry after a 3-day culture in 2D or ULA plates. While the percentage of live cells in 2D were similar between WT and *CDH1* KO cells, the fraction of live cells in ULA were significantly higher in the T47D *CDH1* KO cells (**Fig 5A**), consistent with anoikis resistance. No major differences were observed between the MCF7 WT and *CDH1* KO cells in either condition.

**Figure 5:**
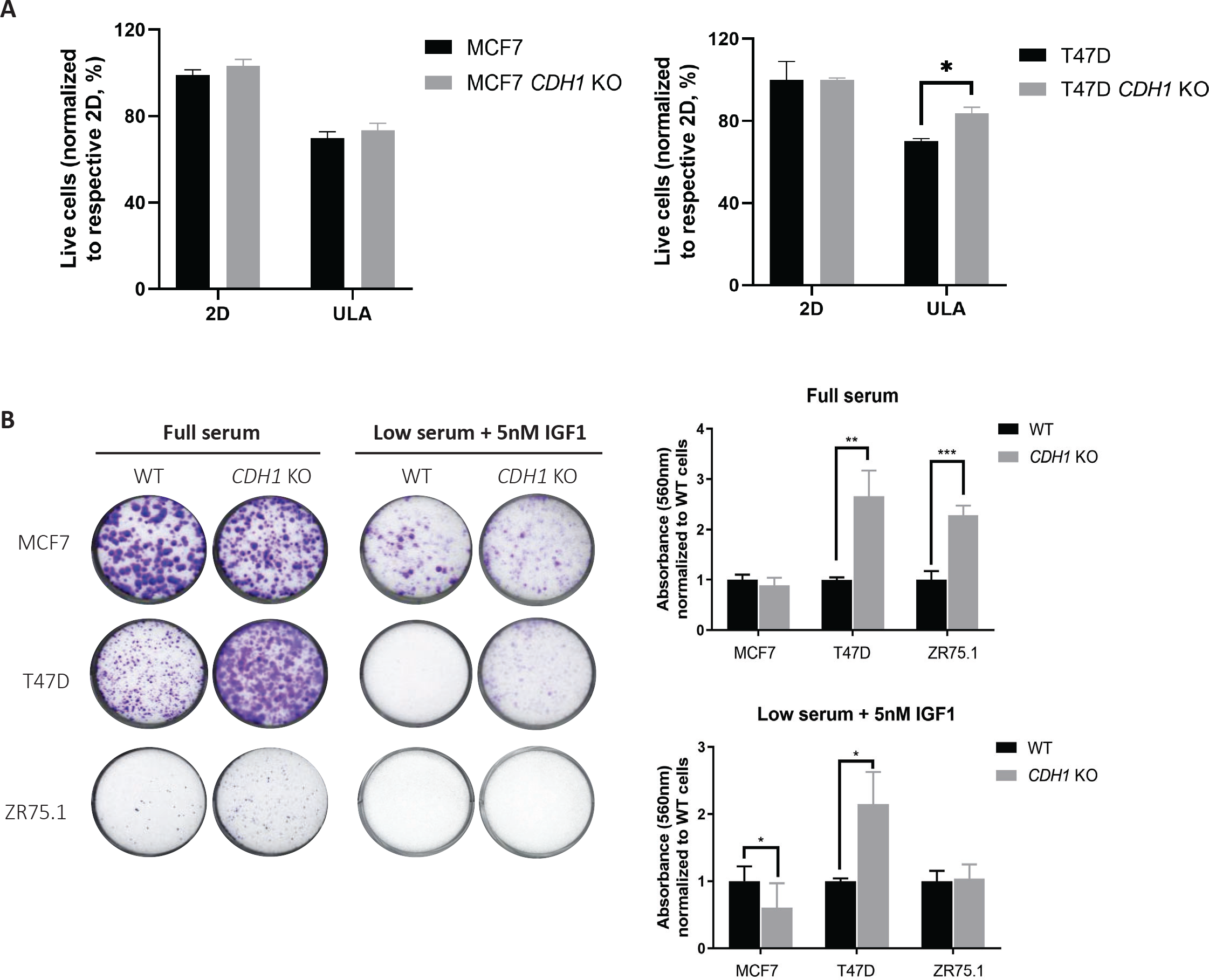
Loss of E-cadherin enhances anoikis resistance and increases cell survival. (A) MCF7 and T47D cells were grown in 2D and ULA plates for 3-4 days and stained with Annexin V and propidium iodide to measure live and apoptotic cells, respectively. Live cell percentage in 2D for each cell line was used to normalize the live cell percentages in ULA plates. Representative experiment shown; N=3 (each with three biological replicates). (B) Cells were plated at 2,000 cells/well in full serum and low serum supplemented with 5nM IGF1 media and stained with 0.5% Crystal Violet after two to three weeks. Representative images are shown. Plates were de-stained with 10% acetic acid, absorbance quantified and graphed after normalization to WT cells of corresponding conditions. Statistical differences were evaluated using two-way ANOVA (*p < 0.05, **p<0.01, ***p<0.001, representative experiment shown, N=3 (each with three biological replicates)).

To further explore cell survival, we performed colony formation assays by plating cells at a low density in either full serum (10% FBS) or low serum (0.5% FBS) containing 5nM IGF1 in 2D plates. T47D *CDH1* KO cells showed increased clonogenic survival in both conditions (**Fig 5B**). *CDH1* KO cells also demonstrated increased colonies in full serum compared to WT, but with no colonies observed in low serum conditions. MCF7 *CDH1* KO cells showed no clear differences in full serum quantifications, however, they formed a greater number of smaller colonies than the WT cells, with similar observations in low serum conditions. Our data support that loss of E-cadherin may play a context-dependent role in anoikis resistance and cell survival.

### Loss of E-cadherin increases collagen I haptotaxis, IGF1-driven migration and serum-driven collagen invasion

We investigated if deletion of *CDH1* in IDC cell lines confers haptotactic migration towards the extracellular matrix (ECM), a phenotype that is pronounced in ILC cell lines (8). We observed a significantly higher migration towards Collagen I by MCF7 and T47D *CDH1* KO cells (**Fig 6A**) compared to their respective WT cells. Multiple studies have reported on E-cadherin loss permitting cell migration and invasion (53–55); conflicting reports on the requirement of E-cadherin for metastasis have also been reported (56). To explore this inconsistency while assessing the importance of IGF1, *CDH1* KO cells were subjected to Transwell migration assays with chemotactic low serum (0.5% FBS) media +/- 5nM IGF1 or full serum (10% FBS) media. All three *CDH1* KO cell models showed increased migration towards serum compared to WT cells **(Fig 6B, C**). Consistent with the IGF1 sensitive nature of these *CDH1* KO cell lines, MCF7 and T47D *CDH1* KO cells also showed significant migration towards IGF1, a phenotype which was successfully halted upon IGF1R/IR inhibitor, BMS-754807 addition (**Supp Fig 4B-D)**.

**Figure 6:**
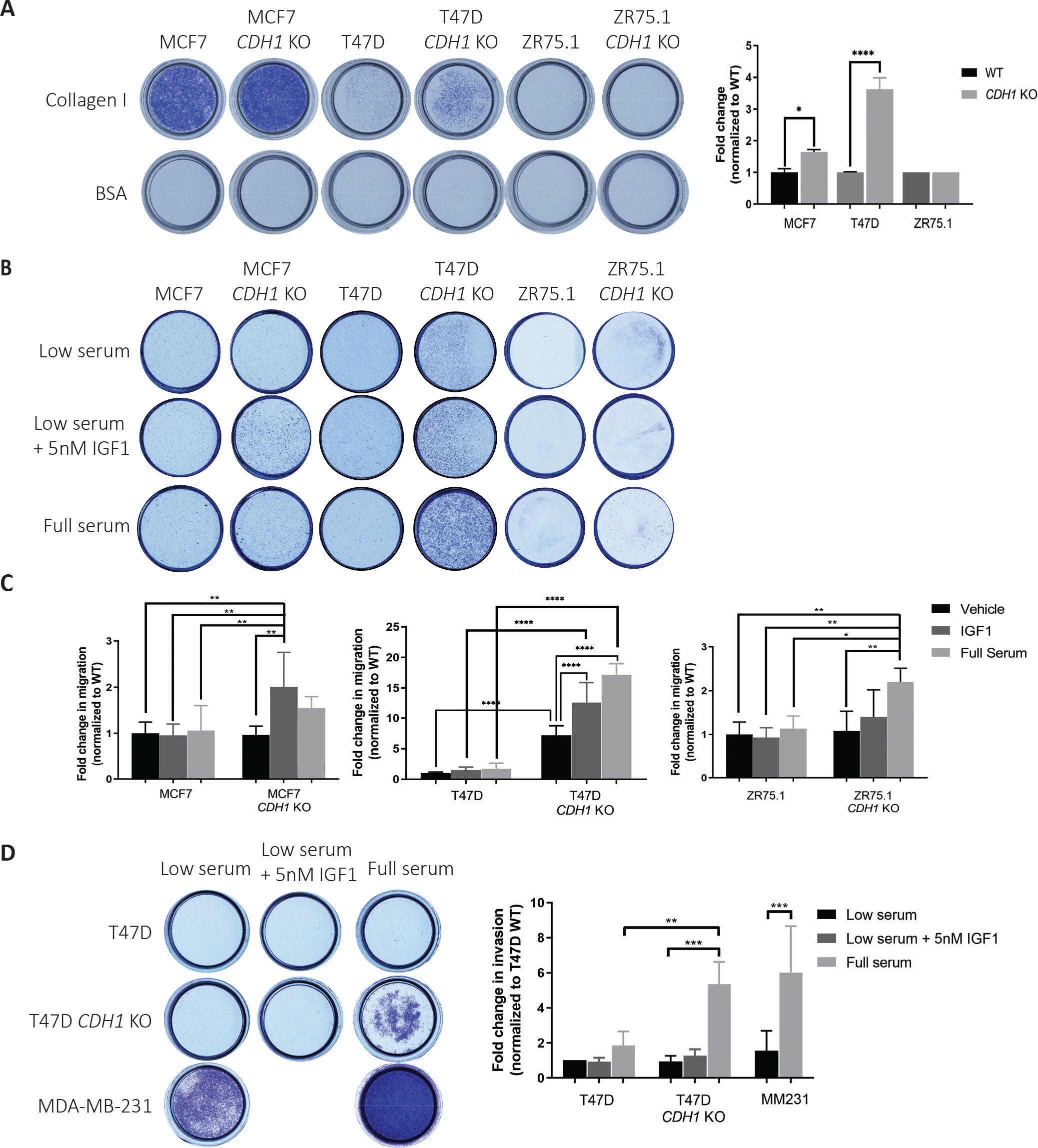
Loss of E-cadherin increases collagen I haptotaxis, IGF1-driven migration and serum-driven collagen invasion. (A) Images and quantification of Crystal Violet-stained Collagen I coated inserts from haptotaxis assays after 72 hours. Migrated cell colonies were quantified with ImageJ and plotted. (B) Images and (C) quantification of Crystal Violet-stained Transwell inserts from migration assays towards the indicated attractants after 72 hours. (D) Images and quantification of Crystal Violet-stained Collagen I inserts from invasion assays towards the indicated attractants after 72 hours. Graphs show representative data normalized to low serum WT cell samples from two to three independent experiments (N=2 biological replicates). p-values from one-way ANOVA statistical testing. *p ≤ 0.05; **p ≤ 0.01; ***p ≤ 0.001; ****p ≤ 0.0001.

Higher migration of T47D *CDH1* KO cells without the presence of any chemotactic gradient highlights the migratory phenotype driven by the independent loss of E-cadherin in this model. Given the haptotactic migration towards Collagen I and migration towards IGF1 and serum, we hypothesized that *CDH1* KO cells may also be able to invade through Collagen I (8) towards a gradient of IGF1 and serum. While there was no Collagen I invasion observed with the MCF7 and ZR75.1 cells (**Supp Fig 4E**), T47D *CDH1* KO cells showed a significant Collagen I invasion towards serum (**Fig 6D**). It is important to note that neither of these three cell lines are invasive by nature and the invasive phenotypes observed in T47D *CDH1* KO cells is promising and requires greater investigation in the context of tumor invasion.

### Loss of E-cadherin sensitizes cells to Fulvestrant and IGF1R/PI3K/Akt/MEK inhibitors in a context dependent manner

Motivated by the increased sensitivity of *CDH1* KO cells to IGF ligands and the potential for clinical translation, we sought to determine if these cells are also sensitive to IGF1R inhibitors. Only T47D *CDH1* KO cells were more sensitive to the tested IGF1R inhibitors compared to WT cells (BMS-754807 and OSI-906) (**Fig 7A, B**). Both MCF7 and ZR75.1 *CDH1* KO cells did not show any significant differences when compared to WT cells (**Supp Fig 5A, B**) although BMS754807 treatment did inhibit IGF1R and Akt signaling in these cell lines, albeit at a lower efficacy in *CDH1* KO cells, potentially owing to their increased ligand sensitivity (**Supp Fig 5C**). Given the higher incidence of *PIK3CA* hotspot mutations and occurrence of *PTEN* loss in ILC (4, 12, 57), we examined if the loss of E-cadherin in these IDC cell lines also sensitizes them to PI3K and Akt inhibitors, Alpelisib and MK-2206, respectively. As with the IGF1R inhibitors, only T47D *CDH1* KO cells showed increased sensitivity to MK2206 and an overall increased sensitivity trend to Alpelisib upon averaging statistical analysis of repeated experiments (**Fig 7C, D, Supp Fig 5A, B, D**). Next, we assessed if the absence of susceptibility t o IGF1R and Akt inhibitors in the MCF7 *CDH1* KO cells might be explained by alternative pathways being activated. We performed combination treatments of BMS-754807 with MEK inhibitor, U0126, and observed a strong additive effect where both MCF7 and T47D *CDH1* KO cells were more sensitive than their corresponding WT cells (**Fig 7E, F**). While this was an expected result in the T47D cells, it is interesting to note the additive effect in the MCF7 *CDH1* KO cells as this suggests the activation of alternative pathways following IGF1R, PI3K and Akt independent inhibitions. We further assessed the synergy effect utilizing the SynergyFinder platform (35), with the ZIP method as output (**Supp Fig. 6A, B**). In MCF7 and T47D WT and *CDH1* KO cells, synergy scores were between -9.717 to 10.995, which validates an additive but not synergistic drug combination effect. As over 90% of ILC tumors are estrogen receptor (ER) positive, we also investigated the combination effects of Akt inhibition with an ER degrader, Fulvestrant, and observed a higher susceptibility in the T47D *CDH1* KO cells (**Fig 7G, Supp Fig 6C**), supporting the notion of targeting the IGF pathway concurrently to targeting ER. These results support the study of combinatory therapies in pre-clinical studies to better target the activation of the IGF pathway following E-cadherin loss. Finally, to analyze pathway inhibition in a more physiologically relevant model, we utilized patient derived ILC and IDC breast organoids (PDO) to compare Akt inhibition sensitivity. As with our cell line results, no significant IDC to ILC comparison was observed, however, ILC organoids did demonstrate a stronger trend in sensitivity to treatment (**Fig 7H, Supp Fig 6D**), strengthening potential translational implications.

**Figure 7:**
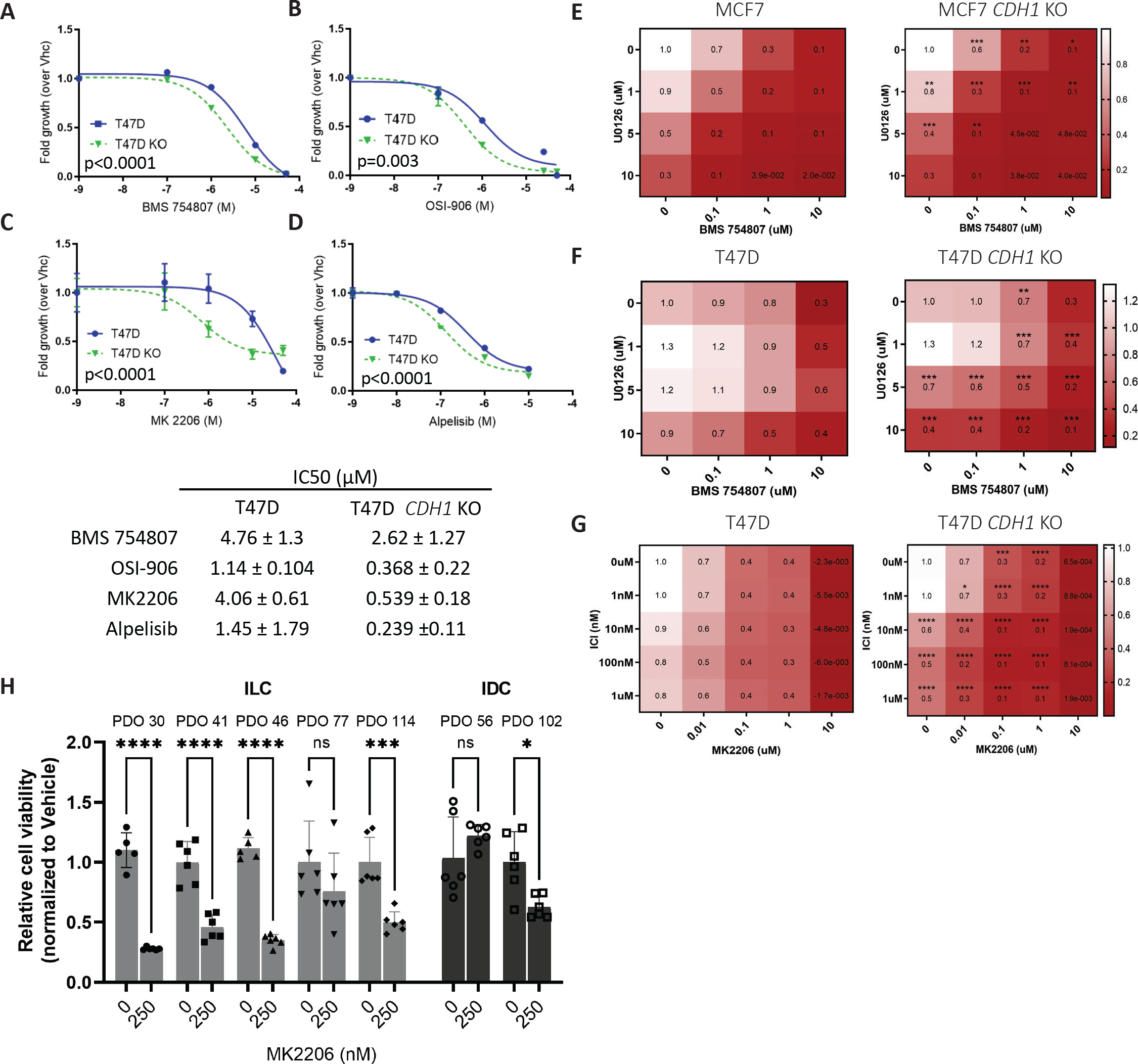
Loss of E-cadherin sensitizes cells to Fulvestrant and IGF1R/PI3K/Akt/MEK inhibitors in a context dependent manner. T47D parental and *CDH1* KO cells were seeded in 96-well 2D plates and treated with IGF1R inhibitors (OSI-906 or BMS-754807), PI3K inhibitor (Alpelisib) or Akt inhibitor (MK2206) for 6 days. Conditions in the panels as follows: (A) BMS- 754807; (B) OSI-906; (C) MK2206 and (D) Alpelisib. CellTiter Glo assay was used to assess cell viability (relative luminescence) and data was normalized to vehicle treated control. (E, F) Cells were treated a combination of MEK inhibitor (U0126) and BMS-754807 for 6 days and viability assessed as above. (G) Cells were treated a combination of Akt inhibitor (MK2206) and Fulvestrant for 6 days and viability assessed as above. (H) Patient derived IDC and ILC organoids were treated with Akt inhibitor (MK2206) for 12 days and viability assessed with CellTiter Glo 3D (relative luminescence), and data was normalized to vehicle treated control. IC50 values for viability were calculated by nonlinear regression and statistical differences evaluated using sum-of-squares Global f-test (p < 0.05; representative experiment shown; N=3 (each with six biological replicates)).

## Discussion

Recent advancements have led to personalized cancer treatments; however, more progress needs to be made to effectively target tumors. Our study highlights the role of E-cadherin in regulating IGF signaling and tumorigenic properties *in vitro*. Genetic deletion of E-cadherin in IDC cells caused reduced cell-cell attachment and morphology transformation into looser cell clusters in suspension. Furthermore, E-cadherin loss was associated with enhanced anchorage independence and anoikis resistance, all of which are features observed in ILC tumors. Additionally, *CDH1* KO cells were more sensitive to IGF1, IGF2 and insulin signaling attributable to an increased receptor availability for ligand binding. This was an IGF-specific phenotype and was not observed with other growth factor signaling pathways tested albeit FGFR4 activations were higher in the *CDH1* KO cells but without significant downstream signaling. Further characterization of *CDH1* KO cells also showed an increased cell survival in clonogenic assays, and enhanced cell migration/invasion in the majority of models, suggesting a role for E-cadherin in controlling metastatic phenotypes. Importantly, a context dependent increase in sensitivity to IGF1R, PI3K, Akt and MEK inhibitors was also observed in our models. This study contributes to an increasing body of work delineating ILC as a unique breast cancer subtype and suggests a potential for targeted therapeutic approaches towards the IGF1R/PI3K/Akt axis owing to the diagnostic loss of E-cadherin in this histological subtype.

Studies have attributed spatial accessibility of IGF1R for ligand binding as a limiting factor in signaling activation (30, 58), and consistent with this notion, we found that loss of E-cadherin confers increased IGF ligand binding by IGF1R, and subsequent pathway activation. Despite previous reports on E-cadherin repressing growth factor receptors (30, 33, 58) and co-localization (58), we were unable to find an association by co-IP despite numerous attempts, suggesting that this is not the mechanism affecting the availability of IGF1R for ligand binding when E-cadherin is present. The formation of a ternary complex involving E-cadherin, IGF1R and integrins is an ongoing topic of investigation thought to affect cell mobility (59), which could additionally complicate immunoprecipitation. Similar interactions have been reported for E-cadherin and EGFR, although we did not observe an increased sensitivity to the EGF ligand in our *CDH1* KO cell lines (30, 60). We, and others, have shown that downregulation of E-cadherin by growth factors can promote EMT (58, 61); others have additionally demonstrated growth factor signaling regulation by E-cadherin, suggesting that a bidirectional feedback loop may exist. These studies were initially almost exclusively performed in the context of EGFR where multiple groups had shown that E-cadherin inhibits ligand-dependent activation of EGFR signaling (30, 62, 63). However, a recent study by Teo and colleagues (13) supported our findings by reporting on ILC cell lines generated from a p53-deficient metastatic mouse model exhibiting had PI3K/AKT pathway activation and enhanced sensitivity to pathway inhibition.

Due to the function of E-cadherin in maintaining cell-cell adhesion, its loss through EMT or genetic deletion is often thought to correlate with increased tumor invasion and metastasis (62, 64). Other studies, however, have suggested a requirement for E-cadherin in metastasis (56).

Here we show that *CDH1* KO cells demonstrate increased migration towards Collagen I (8). The *CDH1* KO cells also showed an enhanced migration towards serum and IGF1, with T47D *CDH1* KO cells showing increased migration even in the absence of a chemoattractant. T47D *CDH1* KO also showed invasion through Collagen I, providing additional *in vitro* evidence supporting that the loss of E-cadherin may enhance metastatic phenotypes. The *in vitro* nature of these findings, however, is an interpretation limitation and *in vivo* experimentation is needed to better understand the role of E-cadherin in metastasis and validate its effect on IGF signaling.

Inhibitors of downstream activators such as PI3K, Akt and MEK have successfully been developed and approved for the clinic. This is paramount in malignancy with a high percentage of mutations leading to *PIK3CA* activation and PTEN loss such as in ILC (4), and maybe more so important given the high IGF pathway activation found in the SCAN-B dataset. Other studies with CRISPR MCF7 *CDH1* KO cells lines have shown that loss of E-cadherin leads to dependency upon ROS1 and enhanced sensitivity to crizotinib (65) for which a clinical trial is currently underway. We report increased sensitivity to IGF1R/PI3K/Akt and MEK inhibitors in T47D *CDH1* KO cells and an increased trend in sensitivity to an Akt inhibitor in ILC PDOs compared to IDC PDOs. Given the genetic landscape of MCF7, T47D and ZR75.1 cells, these models are primed for hyperactive Akt signaling, which has been shown to be further enhanced by loss of E-cadherin. The helical domain mutation in MCF7 cells (E545K) is reported to have a more aggressive phenotype over the kinase domain mutation in T47D (H1047R) (66), which may explain the absence of additional sensitivity in MCF7 *CDH1* KO cells to compounds tested since the *PIK3CA* mutation may effectively mask any potential effect from E-cadherin loss. With the possibility of IGF1R inhibitor efficacy being reduced by hyperactive downstream pathways such as MEK as seen in colon carcinoma (67) and the possibilities of downstream activation inhibition being overhauled by hyperactive receptor tyrosine kinases (68), we tested and observed an additive effect of treating cells with an IGF1R inhibitor in combination with a MEK inhibitor. Further, additive effects of combining Akt inhibitors with Fulvestrant was also found to be beneficial in E-cadherin deficient cells, proving novel therapeutical benefits which require further exploration *in vivo*. Future studies will need to test the efficacy of inhibitors targeting the IGF pathway comparing cells +/- *CDH1 in vivo* and investigate the potential of efficiently exploiting E-cadherin as a biomarker of therapeutic response.

In conclusion, this study delineates the role of E-cadherin in regulating enhanced IGF signaling, controlling metastatic phenotypes, and identifying use of specific tyrosine kinase inhibitors that can effectively target elevated IGF1 signaling activation. We reveal novel findings of increased receptor availability for ligand binding following the loss of E-cadherin and increased susceptibility to IGF1R/PI3K/Akt and MEK inhibitors upon *CDH1* deletion. Our study adds to a growing body of evidence suggesting that *CDH1* may represent a biomarker of response to kinase inhibitors and further exemplifies the need for investigation into the clinical translation of these findings. The *in vitro* nature of our studies in a small panel of cell lines is an important, notable limitation, but notwithstanding, we have robustly revealed important cellular- and contextual-dependent functions of E-cadherin in IGF driven tumor phenotypes. Our findings require translation into *in vivo* models with the goal of validating E-cadherin loss as a biomarker of growth factor receptor inhibitor response in breast cancer as well as the utilization of E-cadherin expression as a vital patient stratification tool in clinical trials.

## Authors’ Contributions

Conception and design: A. Elangovan, S. Oesterreich, J.M. Atkinson, A.V. Lee

Development of methodology: A. Elangovan, J. Hooda, J. Chen, D.D. Brown, P.F. McAuliffe, S. Oesterreich, J.M. Atkinson, A.V. Lee

Acquisition of data: A. Elangovan, J. Hooda, J. Chen, S. Oesterreich, J.M. Atkinson, A.V. Lee

Analysis and interpretation of data: A. Elangovan, J. Hooda, S. Puthanmadhomnarayanan, L. Savariau, M.E. Yates, S. Oesterreich, J.M. Atkinson, A.V. Lee

Writing, review, and/or revision of the manuscript: A. Elangovan, J. Hooda, S. Puthanmadhomnarayanan, L. Savariau, M.E. Yates, J. Chen, D.D. Brown, P.F. McAuliffe, S. Oesterreich, J.M. Atkinson, A.V. Lee

Administrative, technical, or material support: A. Elangovan, J. Hooda, L. Savariau, M.E. Yates, J. Chen, D.D. Brown, P.F. McAuliffe, S. Oesterreich, J.M. Atkinson, A.V. Lee Study supervision: A. Elangovan, S. Oesterreich, J.M. Atkinson, A.V. Lee

## Acknowledgements

The authors would like to thank Osama Shah for data acquisition and bioinformatic input and the Institute for Precision Medicine, a partnership of the University of Pittsburgh and UPMC, for providing the breast cancer patient derived organoids used in these studies. This research was supported in part by the University of Pittsburgh Center for Research Computing through the resources provided.

---

**Supplementary Figure 1.**
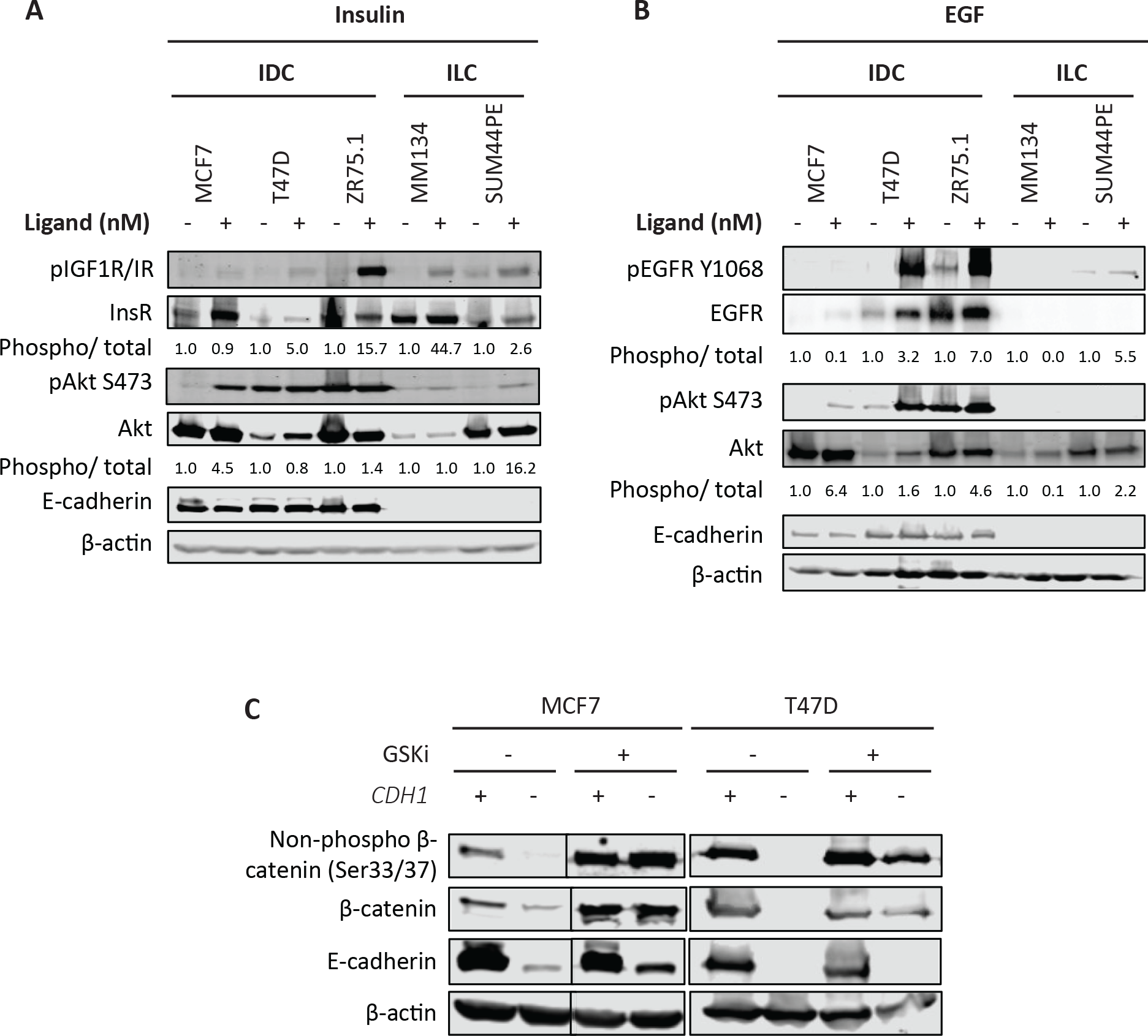
(A) IDC cells (MCF7, T47D, ZR75.1) and ILC cells (MDA-MB-134-VI (MM134) and SUM44PE) were stimulated with doses of insulin or (B) EGF (O-5nM) for 15 minutes following an overnight serum starvation. IGF1R/IR, EGFR and Akt signaling was assessed by Western blotting. Phosphorylated protein levels were normalized over the corresponding total protein levels and loading control β-actin. Ligand treated samples values were further normalized to respective cell line’s vehicle samples. (C) Cells were treated with GSK-3 inhibitor, CHIR 99O21 for 24 hours and harvested for Western blotting. Total and active β-catenin protein levels were assessed. Representative experiment shown for all, n=2-3 for each experiment).

**Supplementary Figure 2.**
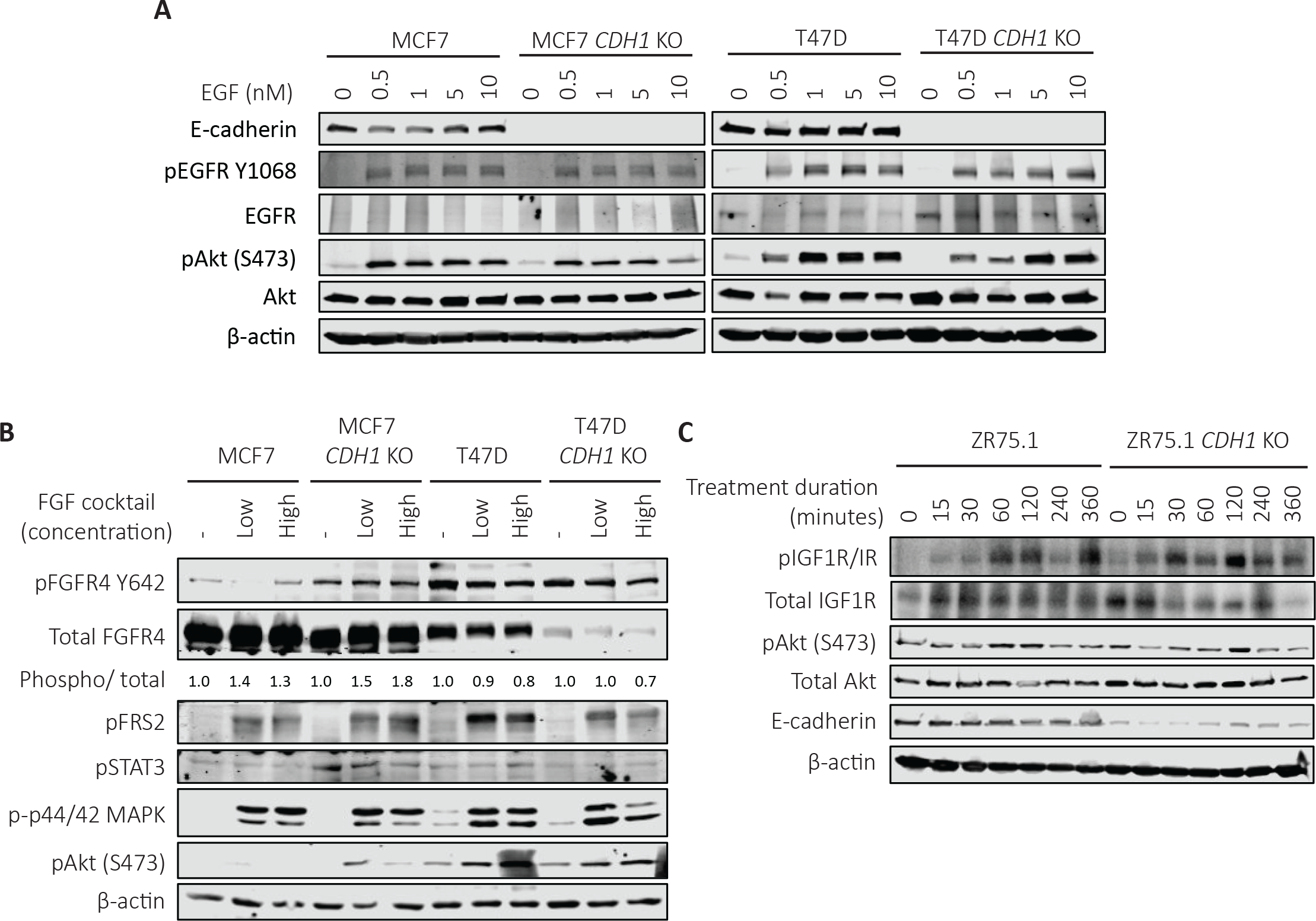
Cells were serum starved overnight and stimulated with (A) EGF (O-1OnM) and (B) cocktail of FGF ligands (1O-5Ong/mL) for 15 minutes. Cells were harvested for Western blot to assess downstream signaling. (C) ZR75.1 parental and *CDH1* KO cells were treated with 1OnM IGF1 for a time course from O-6 hours to assess the duration of signaling activity. Representative experiment shown for all, n=2-3 for each experiment).

**Supplementary Figure 3.**
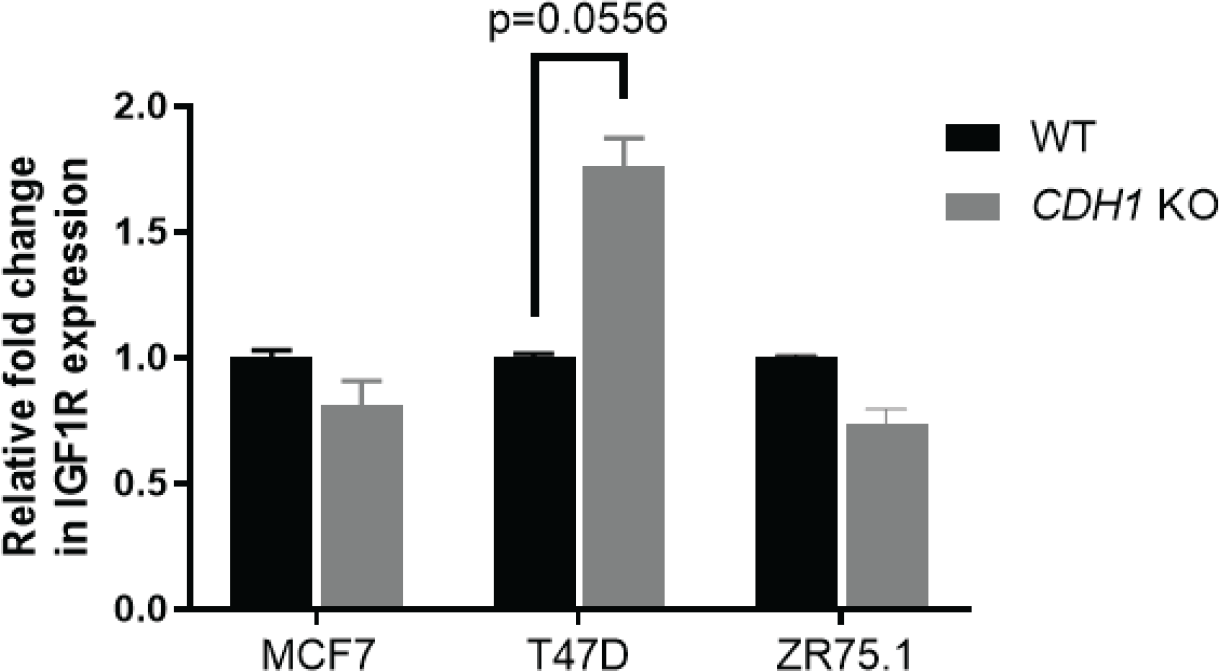
(A) qRT-PCR for IGF1R for the cell line models studied. Fold change in expression were calculated by normalizing delta Ct values to the respective WT values. Statistical differences were evaluated using paired t-test (p=O.O556 for T47D cell pair). Representative experiment shown, n=2 experiments with 2 biological and 3 technical repeats for each cell line.

**Supplementary Figure 4.**
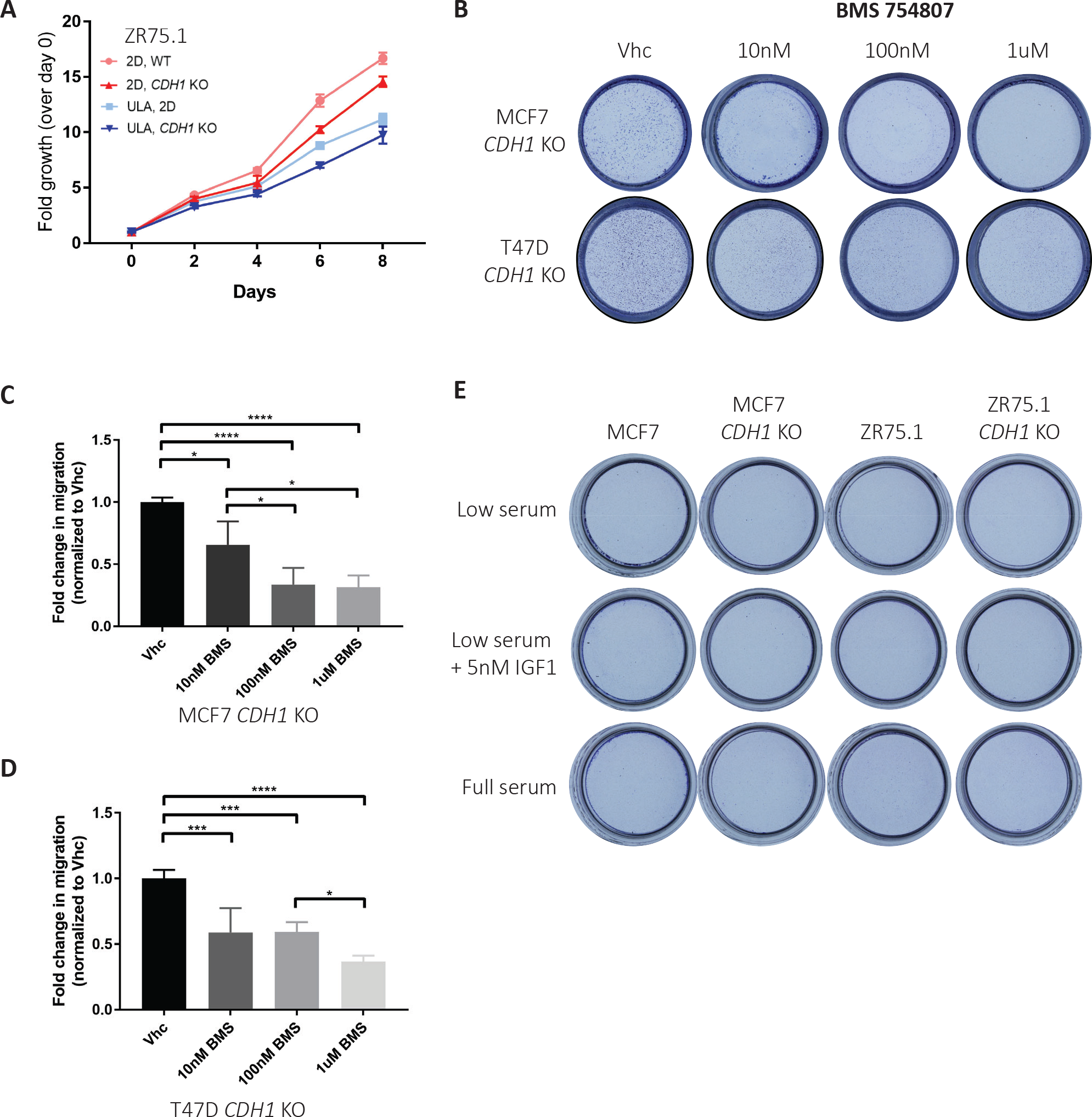
(A) Representative 2D and ULA growth of ZR75.1 parental and *CDH1* KO cells quantified by CellTiter-Glo and normalized to Day O. (B) Images and (C,D) quantification of crystal violet-stained Transwell inserts from migration assays towards low serum + 5nM IGF1 media after 72 hours. Cells were treated with increasing doses of BMS-7548O7 in the upper chamber to assess IGF1 specific migration. Graphs show representative data normalized to vehicle from two to three independent experiments (n=2 biological replicates). p-values are from one-way ANOVA. *p ≤ O.O5; **p ≤ O.O1; ***p ≤ O.OO1; ****p ≤ O.OOO1. (E) Images of crystal violet-stained collagen I inserts from invasion assays towards the indicated attractants after 72 hours.

**Supplementary Figure 5.**
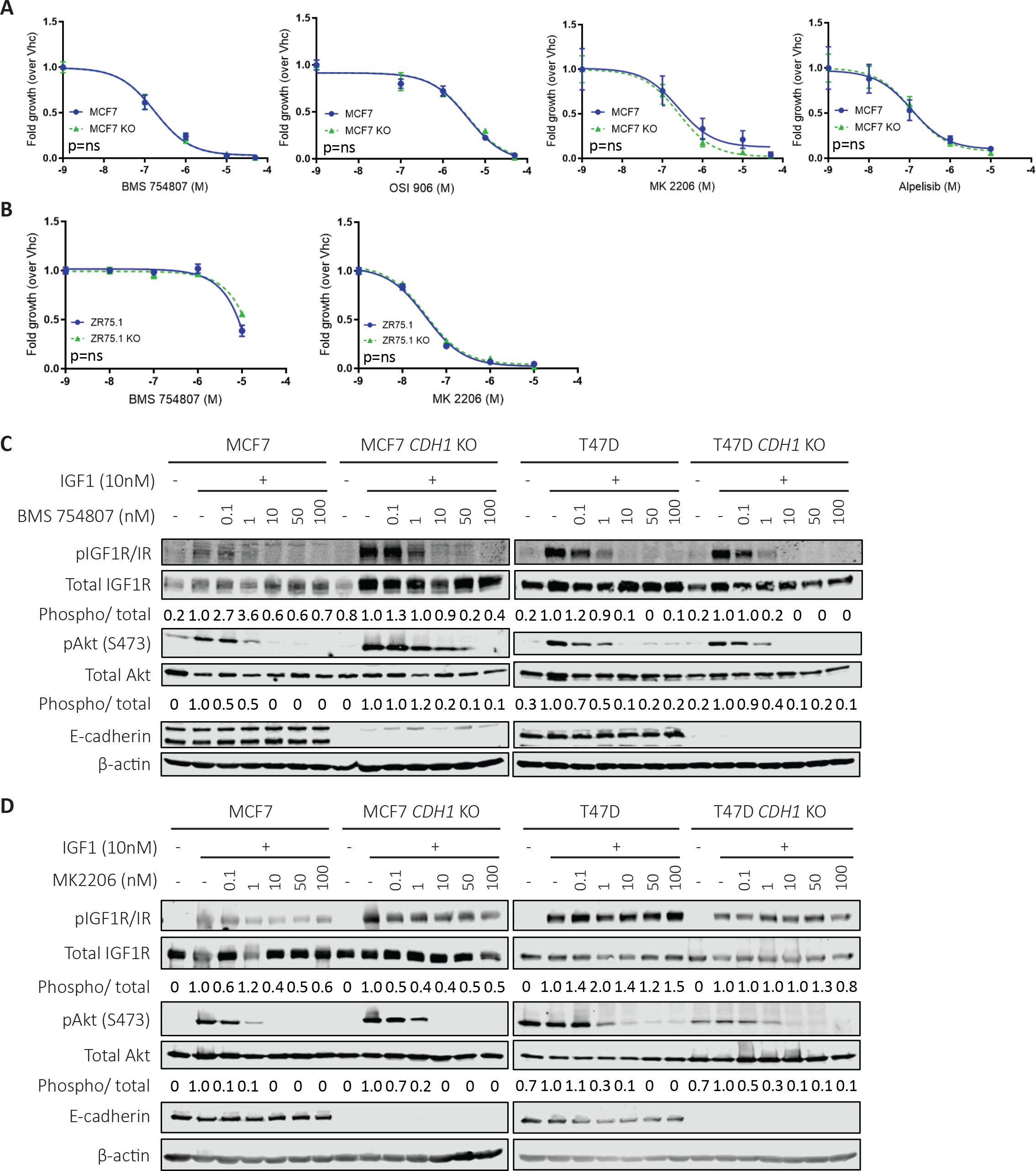
(A) MCF7 parental and *CDH1* KO cells and (B) ZR75.1 parental and *CDH1* KO cells were treated with IGF1R inhibitor (OSI-906 or BMS-754807) or PI3K inhibitor (Alpelisib) or Akt inhibitor (MK2206) for 6 days. CellTiter Glo assay was used to assess cell viability (relative luminescence). IC5O values for viability were calculated by nonlinear regression and statistical differences evaluated using sum-of-squares Global f-test (P < O.O5; representative experiment shown; n=3 each with six biological replicates). Cells were treated with (C) BMS7548O7 and (D) MK22O6 in increasing doses to assess signaling inhibition of the compounds used for cell viability assays. Representative experiment shown, n=2 for each experiment).

**Supplementary Figure 6.**
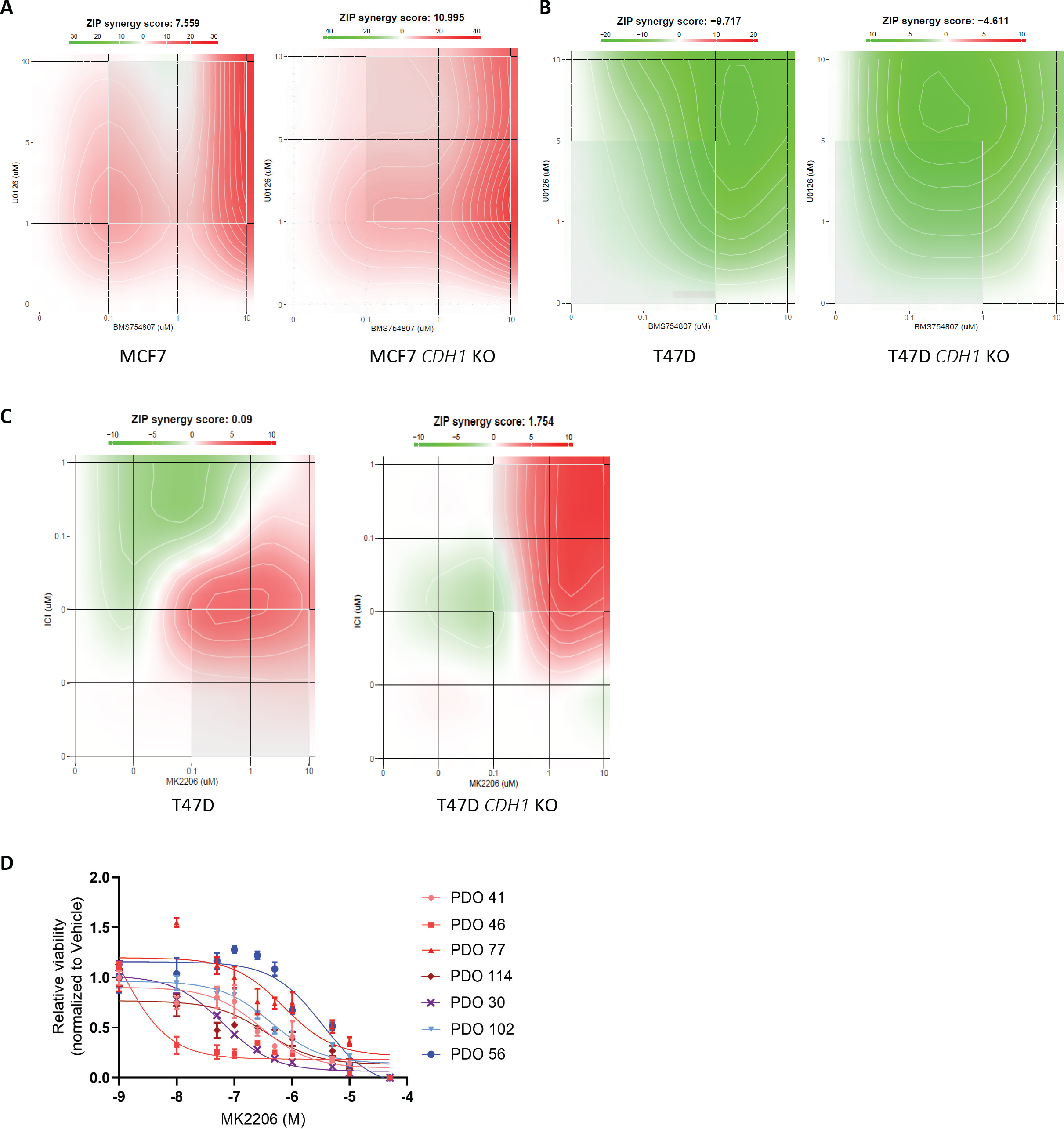
Results from (A) MCF7 WT and *CDH1* KO cells and (B) T47D WT and *CDH1* KO cells were treated with a combination of IGF1R inhibitor (BMS-754807) and MEK inhibitor (U0126) for 6 days. (C) T47D WT and *CDH
1* KO cells were treated with a combination of a SERD (Fulvestrant) and Akt inhibitor (MK22O6) for 6 days. CellTiter Glo assay was used to assess cell viability (relative luminescence). The viability values were input into the SynergyFinder platform and assessed for drug synergy effect. Synergy scores -1O to 1O signify an additive effect of the compounds being tested. (D) Complete dose response analysis of ILC and ILC organoids assayed for sensitivity to MK22O6. Organoids were plated and treated in 96-well for 12 days. CellTiter Glo 3D assay was used to assess cell viability (relative luminescence) and data normalized to vehicle treated control. Representative experiment shown for all, n=2-3 for each experiment).

