## Supplemental Methods for "Loss of E-cadherin Induces IGF1R Activation Revealing a Targetable Pathway in Invasive Lobular Breast Carcinoma"

**Immunoblotting**

Protein for immunoblotting experiments were harvested and analyzed as described previously (35). Membrane blocking was performed with Intercept PBS blocking buffer (LiCOr #927-40000) for one hour at room temperature and probed with primary antibodies overnight at 4°C: pIGF1R/IR (Cell Signaling Technology #3024; RRID:AB_331253), IGF1R (Cell Signaling Technology #3027; RRID:AB_2122378), pAkt S473 (Cell Signaling Technology #4060; RRID:AB_2315049), Akt (Cell Signaling Technology #9272; RRID:AB_329827), InsR (Cell Signaling Technology #3025; RRID:AB_2280448), E-cadherin (BD Biosciences #610182; RRID:AB_397581), β-catenin (BD Biosciences #610154; RRID:AB_397555), p120 catenin (BD Biosciences; #610134; RRID:AB_397537), pEGFR Y1068 (Cell Signaling Technology #2234; RRID:AB_331701), EGFR (Cell Signaling Technology #4267; RRID:AB_2246311), α-Biotin (Cell Signaling Technology #5597; RRID:AB_10828011), non-phospho β-catenin (Cell Signaling Technology #1980; RRID:AB_2650576), pFGFR4 Y642 (Signalway #11836), FGFR4 (Cell Signaling Technology #8562; RRID:AB_10891199), pFRS2 Y196 (Cell Signaling Technology #3864; RRID:AB_2106222), pSTAT3 Y705 (Cell Signaling Technology #9131; RRID:AB_331586) and p-p44/42 MAPK (Cell Signaling Technology #4377; RRID:AB_331775). This was followed by 1 hour room temperature incubation with secondary antibodies (1:10,000; anti-mouse 680LT: LiCor #925-68020; anti-rabbit 800CW: LiCor #925-32211). Membranes were subsequently imaged on the LiCOr Odyssey CLx Imaging system, with band quantifications performed with built in software.

**Immunofluorescence**

Cells were plated at a density of 100,000-200,000 cells/well on glass coverslips (Fisher #12-545-80P) in 24-well plates, fixed on ice in ice cold methanol for 30 minutes and blocked in blocking buffer (0.3% Triton X-100, 5% BSA, 1X DPBS) for 1 hour at room temperature. Primary antibody incubation was performed overnight at 4°C: IGF1R β-subunit (Cell Signaling Technology #3027; RRID: AB_2122378; 1:100), EpCAM (Cell Signaling Technology; #2929; RRID: AB_2098657; 1:100), E-cadherin (Cell Signaling Technology; #3195; RRID: AB_2291471;1:100) and p120 catenin (BD Biosciences; #610134; RRID: AB_397537; 1:100). Secondary antibody incubation was done for 1 hour at room temperature followed by Hoechst 33342 (Thermo Scientific #62249; 1:10000) staining. Coverslips were mounted with Aqua-Poly/Mount (Polysciences #18606-20) and images were taken on a Nikon A1 confocal microscope with a 60X objective.

**Colony formation assay**

Cells were plated at a density of 2000 cells/well in 6-well plates (Fisher #08-772-1B) in either full serum (10% FBS) or low serum (0.5% FBS) with 5nM IGF1 (GroPep Bioreagents #AQU001) supplemented media. Cells were monitored every few days and media refreshed every 4 days. Cells plated in full serum media were fixed with 100% methanol on ice and stained with 0.5% Crystal Violet (Sigma-Aldrich #C0775) in 40% methanol after 14 days while cells grown in low serum, 5nM IGF1 were stained after 21-28 days. Wells were imaged on an Olympus SZX16 dissecting microscope and de-stained with 10% acetic acid in water and quantified by spectrophotometry at 560nm. Statistical differences were tested using two-way ANOVA.

**Co-Immunoprecipitation**

Co-IP was performed with E-cadherin (BD Biosciences #610182; RRID: AB 397581) and IGF1R (Cell Signaling Technology #3027; RRID: AB_2122378) antibodies. Cells were lysed in 20 mM Tris-HCl pH 7.4 with 1% NP-40, 137 mM NaCl and 5 mM EDTA with fresh protease and phosphatase inhibitors (1:100) and quantified using Pierce BCA Protein Assay Kit (Thermo Scientific #23225). 1mg protein from each sample was pre-cleared in 20uL of Pierce™ Protein G Agarose beads (Thermo Fisher Scientific #20398) and incubated in either 3µg of E-cadherin, IGF1R or IgG antibodies (Normal mouse IgG; Millipore #12-371; RRID: AB_145840 and Normal rabbit IgG; Millipore #12-370; RRID: AB_145841) overnight at 4°C with rotation. One 4-hour incubation was performed the next day with 45uL of Pierce™ Protein G Agarose beads at 4°C with rotation. Protein was eluted with Laemmli buffer and analyzed by immunoblotting.

**qRT-PCR**

RNA extraction was performed using RNeasy mini kit (Qiagen #74106) and the RNA quality and amount quantified on NanoDrop. Reverse transcription to cDNA was performed with PrimeScript™ RT Master Mix (Takara Bio #RR036B). RT-PCR was then performed with SsoAdvanced Universal SYBR (Bio-Rad #1726275) with primers as detailed in supplemental materials. Results were normalized to reads from housekeeping gene RPLPO. Statistical differences evaluated using a paired t-test.

**FACS for anoikis resistance**

Cells were stained with APC-Annexin V (BD Biosciences #550474) and Propidium Iodide (BD Biosciences #556463) in 1X Annexin binding buffer (BD Biosciences #556454) for 15 minutes at room temperature. Samples were analyzed on an LSR II Flow cytometer (BD Biosciences) and processed using BD FACSDiva and FlowJo software (BD Biosciences) as previously described (31). Live cell percentages in 2D conditions for each cell line was used to normalize the live cell percentages in ULA conditions. Statistical differences were tested using a two-way ANOVA.
