## Supplemental tables for "Loss of E-cadherin Induces IGF1R Activation Revealing a Targetable Pathway in Invasive Lobular Breast Carcinoma"

Supplemental materials

Table 1: qRT-PCR Primers

| Gene | Forward Sequence (5’ 🡪 3’) | Reverse Sequence (5’ 🡪 3’) |
| --- | --- | --- |
| *CDH1* | GAACAGCACGTACACAGCCCT | GCAGAAGTGTCCCTGTTCCAG |
| *IGF1R* | AGTTATCTCCGGTCTCTGAGG | TCTGTGGACGAACTTATTGGC |

Table 2: Antibodies Used in IB, IF and IHC Experiments

| Protein | Company | Catalog Number | Host Species | Dilution |
| --- | --- | --- | --- | --- |
| β-Actin | Sigma | A5441 | Mouse | 1:10000 |
| AKT | Cell Signaling | 9272 | Mouse | 1:1000 |
| pAKT^S473^ | Cell Signaling | 4060 | Rabbit | 1:1000 |
| E-cadherin | BD Biosciences | 610182 | Mouse | 1:1000 |
| E-cadherin | Cell Signaling | 3195 | Rabbit | 1:100 |
| pIGF1R/ InsR | Cell Signaling | 3024 | Rabbit | 1:500 |
| pEGFR Tyr1068 | Cell Signaling | 2234 | Rabbit | 1:1000 |
| IGFR | Cell Signaling | 3027 | Rabbit | 1:1000 |
| InsR | Cell Signaling | 3025 | Rabbit | 1:1000 |
| EGFR | Cell Signaling | 4267 | Rabbit | 1:1000 |
| IRS1 | Cell Signaling | 2390 | Rabbit | 1:1000 |
| B-catenin | BD Biosciences | 610154 | Mouse | 1:1000 |
| p120 | BD Biosciences | 610134 | Mouse | 1:200 |
| a-Biotin | Cell Signaling | 5597S | Rabbit | 1:1000 |
| Non-phospho B-catenin (Ser45) | Cell Signaling | 19807T | Rabbit | 1:1000 |
| pFGFR4 Y642 | Signalway Antibody | 11836 | Rabbit | 1:1000 |
| FGFR4 | Cell Signaling | 8562 | Rabbit | 1:1000 |
| pFRS2 Tyr196 | Cell Signaling | 3864 | Rabbit | 1:1000 |
| pSTAT3 Tyr705 | Cell Signaling | 9131 | Rabbit | 1:1000 |
| P-p44/42 MAPK | Cell Signaling | 4377 | Rabbit | 1:1000 |
| EpCAM | Cell Signaling | 2929 | Mouse | 1:100 |
| Anti-Mouse AlexaFluor488 | Invitrogen | A11017 | Mouse | 1:200 |
| Anti-Rabbit AlexaFluor488 | Invitrogen | A21206 | Rabbit | 1:200 |
| Anti-Rabbit AlexaFluor647 | Invitrogen | A27040 | Rabbit | 1:200 |

Table 3: Primers for *CDH1* deletion Sanger Sequencing

| Gene | Forward Sequence (5’ 🡪 3’) | Reverse Sequence (5’ 🡪 3’) |
| --- | --- | --- |
| *CDH1* | AGGAGACTGAAAGGGAACGGTG | GTGCCCTCAACCTCCTCTTCTT |
